# Dietary long-chain omega-3 fatty acids are related to impulse control and anterior cingulate function in adolescents

**DOI:** 10.1101/379263

**Authors:** Valerie L. Darcey, Goldie A. McQuaid, Diana H. Fishbein, John W. VanMeter

## Abstract

Impulse control, an emergent function modulated by the prefrontal cortex (PFC), helps to dampen risky behaviors during adolescence. Influences on PFC maturation during this period may contribute to variations in impulse control. Availability of omega-3 fatty acids, an essential dietary nutrient integral to neuronal structure and function, may be one such influence. This study examined whether intake of energy-adjusted long-chain omega-3 fatty acids (eicosapentaenoic acid (EPA) + docosahexaenoic acid (DHA)) was related to variation in impulse control and PFC activity during performance of an inhibitory task in adolescents (n= 87; 51.7% female, mean age 13.3+1.1 years) enrolled in a longitudinal neuroimaging study. Intake of DHA+EPA was assessed using a food frequency questionnaire and adjusted for total energy intake. Inhibitory control was assessed using caregiver rating scale (BRIEF Inhibit subscale) and task performance (false alarm rate) on a Go/No-Go task performed during functional MRI. Reported intake of long-chain omega-3 was positively associated with caregiver ratings of adolescent ability to control impulses (p=0.017) and there was a trend for an association between intake and task-based impulse control (p=0.072). Furthermore, a regression of BOLD response within PFC during successful impulse control (Correct No-Go versus Incorrect NoGo) with energy-adjusted DHA+EPA intake revealed that adolescents reporting lower intakes display greater activation in the dorsal anterior cingulate, potentially suggestive of a possible lag in cortical development. The present results suggest that dietary omega-3 fatty acids are related to development of both impulse control and function of the dorsal anterior cingulate gyrus in normative adolescent development. Insufficiency of dietary omega-3 fatty acids during this developmental period may be a factor which hinders development of behavioral control.

## Introduction

Development of cortical gray matter follows a regionally-specific, nonlinear maturation pattern, whereby gray matter volume generally increases in childhood, peaks in late childhood/early adolescence and declines into young adulthood (Giedd et al., 1999; Gogtay et al., 2004; Lenroot & Giedd, 2006). Gray matter thinning during adolescence is thought to reflect, at least in part, synaptic pruning and refinement (Huttenlocher & Dabholkar, 1997), and is associated with improvements in cognitive function and behavior (Casey, Giedd, & Thomas, 2000). Within the PFC, these dynamic developmental processes occur rapidly during the adolescent years and are thought to underlie improvements in executive function, including impulse control (Rubia et al., 2000; Tamm, Menon, & Reiss, 2002). Response inhibition, an individual’s ability to inhibit his/her actions, is one executive function that is integral to developing the ability to delay gratification (Steinbeis, Haushofer, Fehr, & Singer, 2014) – a cornerstone of long-term achievements (Mischel, Shoda, & Rodriguez, 1989). Response inhibition improves from childhood, through adolescence, and into young adulthood (Luna, Padmanabhan, & O’Hearn, 2010). This maturational process is supported by regionally specific changes in activation within the PFC (Ordaz, Foran, Velanova, & Luna, 2013; Rubia et al., 2006) and the protracted development of the PFC may reflect a period of vulnerability to various environmental and biological factors.

Omega-3 fatty acids are a class of long-chain polyunsaturated fats that can only be obtained via diet. Docosahexaenoic acid (DHA), an omega-3 fatty acid (hereafter, N3) found in marine sources, is the only fatty acid of its class relevant to the central nervous system (Stillwell & Wassall, 2003). Variation in dietary DHA is reflected in variation of DHA content of phospholipids measured both in the peripheral tissues and in the central nervous system (Connor, Neuringer, & Lin, 1990; Hulbert, Turner, Storlien, & Else, 2005; Moriguchi & Salem, 2003). Within the central nervous system, the distribution of DHA is particularly concentrated in the neuronal membranes of the PFC (Bradbury, 2011), highlighting its importance in a region that is critical for executive function. DHA accrues rapidly in the PFC from the perinatal period though the first 18 years of life, with little increase in PFC DHA content after the second decade of life (Carver, Benford, Han, & Cantor, 2001) suggesting that the adolescent years are a crucial time to ensure adequate accrual of DHA in the PFC.

DHA plays a key role in neuronal functions critical to development. DHA promotes membrane fluidity and the interaction of embedded proteins (Stillwell & Wassall, 2003), neuronal signaling and the resolution of inflammation (McNamara & Carlson, 2006; Mitchell, Gawrisch, Litman, & Salem, 1998), is associated with greater neuronal size (Ahmad et al., 2002), enhances dendritic spine density thereby promoting synaptic formation (Wurtman, Cansev, & Ulus, 2009), and facilitates cortical pruning (de Velasco et al., 2012). Low levels of omega-3 fatty acids in the diet reduce DHA incorporation in synaptic membranes (Hulbert et al., 2005), which may be detrimental to development of the PFC and inhibitory control. In adults, omega-3 fatty acid status is associated with greater anterior cingulate cortex (ACC) thickness (Zamroziewicz, Paul, Rubin, & Barbey, 2015) and volume (Conklin, Gianaros, et al., 2007). In children, blood DHA levels were related to PFC activity during sustained attention, activity which is enhanced with supplementation, including within the ACC (McNamara et al., 2010). Compared to boys with high blood DHA, boys with low DHA displayed reduced functional connectivity in cortical attention networks, including the right ACC, during sustained attention (Almeida, Jandacek, Weber, & McNamara, 2016). Low blood levels of DHA are associated with Attention Deficit Hyperactivity Disorder in children (Burgess, Stevens, Zhang, & Peck, 2000; Chen, Hsu, Hsu, Hwang, & Yang, 2004; Stevens et al., 1995) and self-reported impulsivity in adults (Conklin, Harris, et al., 2007). Moreover, low intake of dietary sources of omega-3 fatty acids produces impulsive behaviors in animal models (Levant, Zarcone, & Fowler, 2010) and is associated with externalizing behaviors in children (Gispert-llaurado et al., 2016). Supplementation has been shown to improve impulse control task performance in adults (Fontani et al., 2005). These studies suggest that N3 fatty acids may be important for prefrontal structure and function, particularly in the anterior cingulate cortex.

Dietary profile of polyunsaturated fat intake by Americans has changed dramatically over the last century, resulting in a net decrease in effective dietary omega-3 fatty acids (Blasbalg, Hibbeln, Ramsden, Majchrzak, & Rawlings, 2011) and adolescent diets have been found to be poor in sources of N3 fatty acids (Cutler, Flood, Hannan, & Neumark-Sztainer, 2009). Decreased intake of N3 fatty acids among adolescents may be of particular concern given that DHA rapidly accumulates in membranes of PFC gray matter primarily during the first two decades of life (Carver et al., 2001). Low intake during a critical window of DHA accrual in a brain region undergoing major dynamic development has the potential to negatively impact cortical function and related developing behaviors such as impulse control.

In the present study, we investigated the relationship between intake of long chain omega-3 fatty acids and prefrontal function during impulse control in a cross-sectional sample of typically developing adolescents. Adolescent participants completed a food frequency questionnaire from which an energy-adjusted Omega-3 Index was computed. This Index was then related to prefrontal activity and task performance during a Go/No-Go task while undergoing fMRI, as well as caregiver-rated ability of the adolescent to inhibit impulses. Given the evidence reviewed above, we expected that greater intake of energy-adjusted Omega-3 Index would facilitate task performance. Specifically, we hypothesized that higher levels of Omega-3 Index would be associated with lower PFC activity (i.e., greater efficiency) and increased ability to inhibit prepotent responses (i.e., correct No-Go’s). We also predicted higher levels of Omega-3 Index would be associated with better inhibitory behavior as rated by caregivers.

## Methods

Participants were recruited as a part of a longitudinal neuroimaging study, the Adolescent Development Study (ADS), aimed at identifying neurobiological precursors and consequences of early drug and alcohol initiation and escalation. Full details of the methods are described elsewhere (Fishbein, Rose, Darcey, Belcher, & VanMeter, 2016). In brief, adolescents in a narrow age range (11-13 years old) were recruited and the main exclusionary criteria included prior substance use, left-handedness, conditions rendering MRI unsafe, history of head trauma, and neurodevelopmental disorders. Participants taking psychostimulant (centrally acting) medications were permitted to enroll in the study if study visits could be scheduled during normally occurring medication “holidays”. Demographic, neurocognitive, drug and alcohol use surveys, and imaging assessments were conducted at baseline and repeated 18- and then 36-months later. The data reported here were collected during the baseline and 18-month visits. The Georgetown Institutional Review Board approved all study procedures and adolescents and their caregivers provided assent and consent prior to all data collection.

### Food frequency questionnaire

Adolescents completed a paper-based food frequency questionnaire called the Harvard Youth/Adolescent Food Frequency Questionnaire (YAQ) to assess usual diet over the past year. The YAQ is a widely-used, scantron questionnaire validated for ages 9-18, which provides a dietary analysis based on a retrospective assessment of usual frequency and portions of 152 food items consumed over the past 12 months (Rockett et al., 1997). Questionnaires were completed at the end of study visits. Participants were paid $15 via Amazon gift card for completing the 30-minute survey. Nutrient output was compiled using 2011 nutrient tables (Rockett et al., 1997).

While evidence suggests that DHA is the most relevant long chain omega-3 fatty acid, there have been reports of its long chain precursor, eicosapentaenoic acid (EPA) also being related to neural and cognitive outcomes (Bauer et al., 2014; Bauer, Crewther, Pipingas, Sellick, & Crewther, 2013). Thus, our analyses are based on the reported intake of DHA + EPA, known as the Omega-3 Index (Harris & Von Schacky, 2004), adjusted for total energy consumed [(EPA grams + DHA grams) / total calories] (Subar et al., 2001) and then scaled by 1000 calories to represent long chain N3 fatty acid consumption per 1000 calories of intake (hereafter, energy-adjusted N3 Index). Reported intake of DHA is adjusted for total energy intake to reduce extraneous variation (diets higher in total calories may also be higher in fats consumed) (Willett, Howe, & Kushi, 1997). A square root transformation was applied to the energy-adjusted N3 Index in order to minimize influence of a few participants reporting highest omega-3 intakes. A subsample of adolescents participated in a validation sample, providing both YAQ responses and blood samples for whole blood essential fatty acid analysis (n=19). Energy-adjusted dietary Omega-3 index and Omega-3 index observed in blood (EPA+DHA) were highly correlated (*r_s_*=0.660, p=0.002), similar to other studies (e.g., Almeida et al., 2017; Dahl, Maeland, & Bjorkkjaer, 2011; Marangoni, Colombo, Martiello, Negri, & Galli, 2007).

### Additional adolescent measures

Participants’ intelligence was assessed using a developmentally appropriate battery (Kauffman Brief Intelligence Questionnaire; K-BIT) (Kaufman & Kaufman, 1990). To account for potential differences in physical maturation, adolescents completed the Pubertal Development Scale (Carskadon & Acebo, 1993; Petersen et al., 1988) consisting of a series of questions about progress of physical development, asking respondents to evaluate the degree to which a specific physical change (such as skin/voice changes, growth spurt, breast development, and facial hair) has occurred. During study visits, body mass index (BMI) (kg/m^2^) sex and age-specific z-scores and percentiles were calculated using weight measured with a digital scale (Health-O-Meter Professional 394KLX) and height measured via stadiometer (SECA 216 Wall-mount Mechanical measuring rod; triplicate measures within 0.5 cm, averaged) applied to 2000 CDC Growth Charts (Kuczmarski et al., 2000).

### Family socioeconomic status

Caregivers of participants were interviewed to collect information on parental education and income to calculate an index of household socioeconomic status (SES index) using a method adapted from Manuck and colleagues (2010). Maternal and paternal cumulative years of education were averaged and a standardized education z-score was calculated for each participant. Reported level of total annual household income prior to taxes (ranging from less than $5,000 to greater than or equal to $200,000) was converted to standardized income z-score for each participant. SES index was computed by averaging standardized values for the income and education variables for each participant, then subsequently re-standardizing to achieve a distribution with a 0-centered mean and standard deviation of 1 for the full sample (N=135).

## ASSESSMENTS OF RESPONSE INHIBITION

### Behavior Rating Inventory of Executive Function (BRIEF)

An 86-item psychometrically validated questionnaire (Gioia, Isquith, Guy, & Kenworthy, 2000) to assess facets of executive abilities was administered to primary caregivers of participants. Caregivers rated the frequency that their child’s behaviors were problematic as “never”, “sometimes”, or “often”. The questionnaire yields 8 non-overlapping scales, of which the Inhibit subscale, reflecting the ability to control impulses or stop behavior, was of interest to the current study. Higher scores suggest higher level of dysfunctional behavior. Normative values for age and sex (t-scores) are reported here. No responses were classified as “inconsistent” and all were included in the analysis.

### Go/No-Go task

Adolescents completed a simple Go/No-Go functional MRI task, which elicits neuronal activity related to response inhibition (Menon, Adleman, White, Glover, & Reiss, 2001; Simmonds, Pekar, & Mostofsky, 2008). This task uses a hybrid design with alternating blocks of event-related Go/No-Go (45 seconds) and Fixation (12-16 seconds) each repeated 5 times (total time: 5:02 minutes). During the Go/No-Go blocks, a series of 30 letters is presented for 200 ms each, followed by a 1300 ms fixation. Subjects are instructed to press the button held by their right hand as quickly as possible for every letter (“Go” trials) except the letter ‘Q’ (“No-Go” trials). A total of 150 trials are presented in this design of which 18% are No-Go trials. The task was implemented in E-prime and completed during MR imaging.

While relative accuracy of performance on the Go/No-Go is an indication of attention, errors of commission (response to the target when the correct action is a withheld response) are an indication of poor response inhibition (Riccio, Reynolds, Lowe, & Moore, 2002). Thus, we analyzed the percentage of incorrect No-Go’s (also known as false alarms), and reaction time to Go trials (reflecting processing speed). Responses faster than 150 ms were excluded from behavioral analysis and not modeled in the fMRI analysis (below) to minimize analysis of anticipatory responses (i.e., unlikely to reflect true stimulus processing). Hit rate, or response rate to correct Go (limited to responses longer than 150 ms), was used to determine whether participants were adequately engaged with the task and a hit rate of at least 70% of Go trials was required for inclusion in the analyses.

## MRI PROTOCOL

### Data acquisition

MRI scans were performed on a 3 T scanner (Siemens Tim Trio) at the Center for Functional and Molecular Imaging, Georgetown University (Washington, DC) using a 12-channel radio frequency head coil. For anatomical localization and spatial normalization a structural MRI acquisition was collected using a 3D T1-weighted MPRAGE image with the following parameters: TR/TE=1900/2.52 ms, TI=900 ms, 176 slices and slice resolution= 1.0 mm^3^. FMRI acquisition used T2*-weighted gradient-echo planar imaging (EPI). The blood oxygenation level dependent (BOLD) functional MRI acquisition parameters were: TR/TE 2500/30 ms, 90° flip angle, in-plane resolution 3.0 mm^2^, 47 slices and slice thickness=3.0 mm. Data for the Go/No-Go task were collected in one run.

### fMRI Data preprocessing

Image processing and statistical analysis was carried out using SPM8 (http://www.fil.ion.ucl.ac.uk/spm) including correction for sequential slice timing and realignment of all the images to the mean fMRI image to correct for head motion artifacts between images. Realigned images were then co-registered with the anatomical MPRAGE. The MPRAGE was segmented and transformed into the Montreal Neurological Institute (MNI) standard stereotactic space using affine regularization and nonlinear registration with tissue probability maps. Lastly, these transformation parameters were applied to normalize the fMRI images into MNI space, after which the data were spatially smoothed using a Gaussian kernel of 6 mm^3^ full-width half maximum (FWHM). A scrubbing algorithm utilizing framewise displacement (FD) was used to assess participant movement during the fMRI scans (Power, Barnes, Snyder, Schlaggar, & Petersen, 2012). Participants were excluded from analyses if they had more than 1 mm frame-wise displacement in over 20% of their volumes during the fMRI scans.

### First-level analysis

Regressors for the first-level analysis included vectors coding for correct Go and No-Go and incorrect Go and No-Go trials. Six regressors of no interest from the motion correction parameters were included to minimize signal changes related to head movement. The fMRI responses were convolved with the canonical hemodynamic response function and a 128-s temporal high-pass filter was applied to the data to exclude low-frequency artifacts such as MRI signal drift. The contrast of interest was successful inhibitions to examine activation associated with cognitive control of behavior (i.e., Correct No-Go > Incorrect No-Go).

### fMRI statistical analysis

First level contrasts were entered into a linear regression with the square root transformed energy-adjusted N3 Index. Given the relative importance of DHA to PFC function, second-level group analyses were constrained to a specific search region with an explicit mask of bilateral frontal lobe gray matter, created using AAL, Wake Forest Pick Atlas (Maldjian, Laurienti, Kraft, & Burdette, 2003). Clusters were defined in SPM8 using an uncorrected threshold of p = 0.001 with a cluster extent of 10 voxels and were determined to survive correction for multiple comparison at the cluster level using FWE of p <0.05. Peak MNI coordinates were extracted from each surviving cluster and anatomical descriptions were identified using the AAL atlas.

### Data reduction/exclusions

Of 135 participants enrolled in the parent study, 126 completed the food frequency questionnaire. Seven of these responses were excluded from analysis for the following reasons: one participant for implausibly high caloric intake (>13,000 kcal per day (Cutler et al., 2009)); three for leaving excessive numbers of questions blank; three due to factors revealed post-enrollment that may affect neurodevelopment (n=2 reporting drug/alcohol prior to study enrollment, n=1 Tourette’s syndrome diagnosis).

Of 119 participants with eligible food frequency questionnaire data, thirteen were excluded from the behavioral analysis (three due to technical data collection errors; ten for hit rates < 70%). Of 106 participants remaining with eligible diet surveys and Go/No-Go behavior, 19 were excluded on the basis of imaging data (4 due to braces, 3 were missing full imaging data, 2 were missing part of the brain image, and 10 for excessive motion). Final analyses were restricted to participants with eligible imaging, behavioral and dietary data (n=87).

### Statistical analyses

Data analysis was conducted in IBM SPSS Statistics 24 (IBM Corp. Released 2016. IBM SPSS Statistics for Windows, Version 22.0. Armonk, NY: IBM Corp.). The distributions of dependent variables were confirmed via Shapiro-Wilk test. The BRIEF Inhibit t-score, false alarms, and processing speed possessed skewed/kurtotic distributions, as indicated by *Spearman’s Rho* (*r_s_*). Non-parametric analyses were therefore used for these three DVs. The distribution of the independent variable, energy-adjusted N3 Index, was transformed by taking the square root in order to minimize influence of a few participants reporting high N3 intakes. Multiple linear regressions were calculated using rank-transformed data to examine the relative contribution of energy-adjusted N3 Index and demographic predictors (i.e., age, socioeconomic status index) on behavioral outcomes for significant correlations.

## Results

Participant characteristics are presented in Table 1. Energy-adjusted N3 Index was unrelated to age (*r*=0.182; p=0.091; n=87), SES index z-score (*r_s_*=0.089, p=0.420, n=84), pubertal development score (*r*=0.044, p=0.686, n=87), BMI z-score (*r*=−0.072, p=0.505, n=87) or KBIT IQ score (*r_s_*=0.150, p=0.174, n=84). Males and females reported similar intakes of N3 Index (Mann-Whitney-U test p=0.363).

**Table 1.**
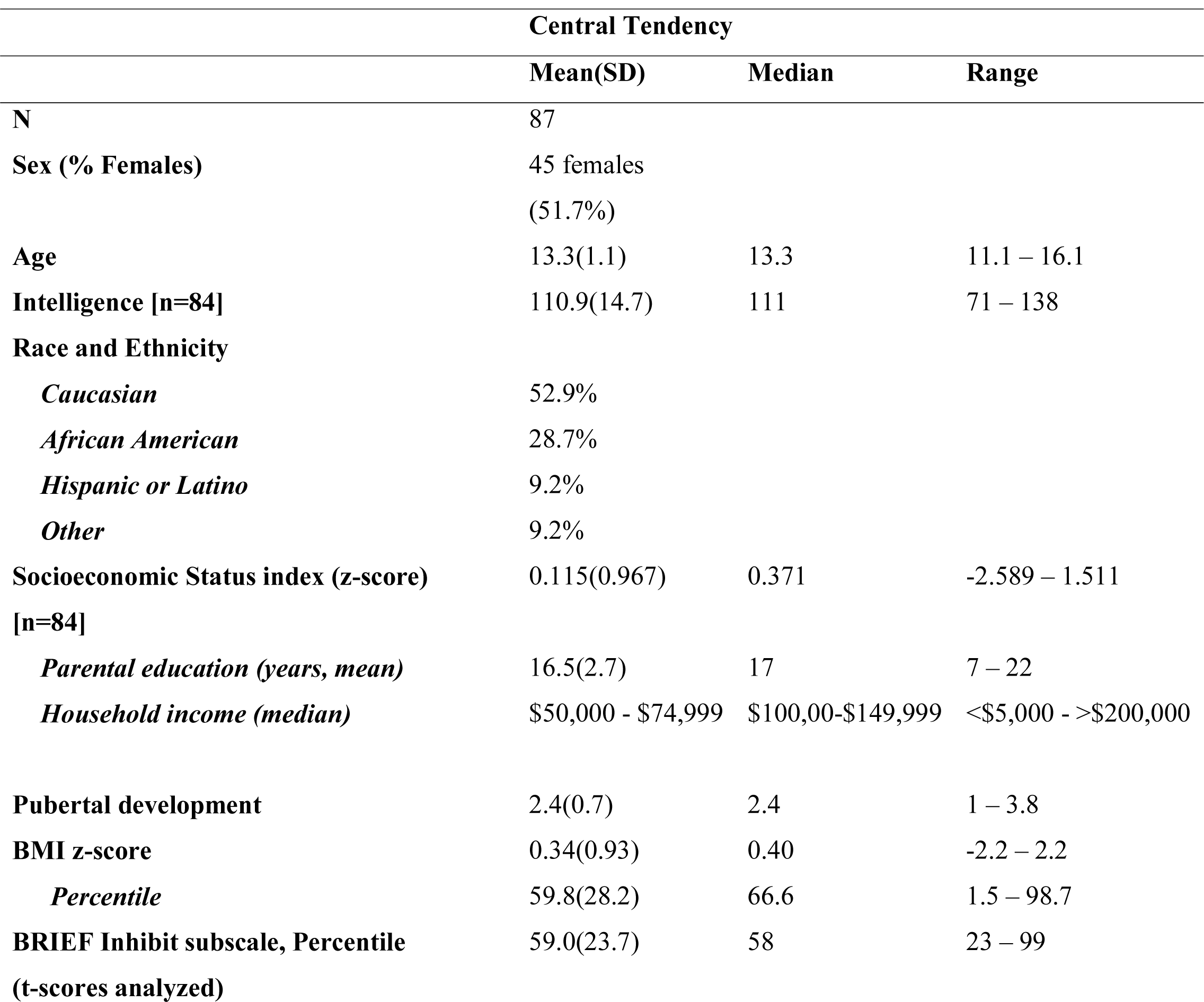
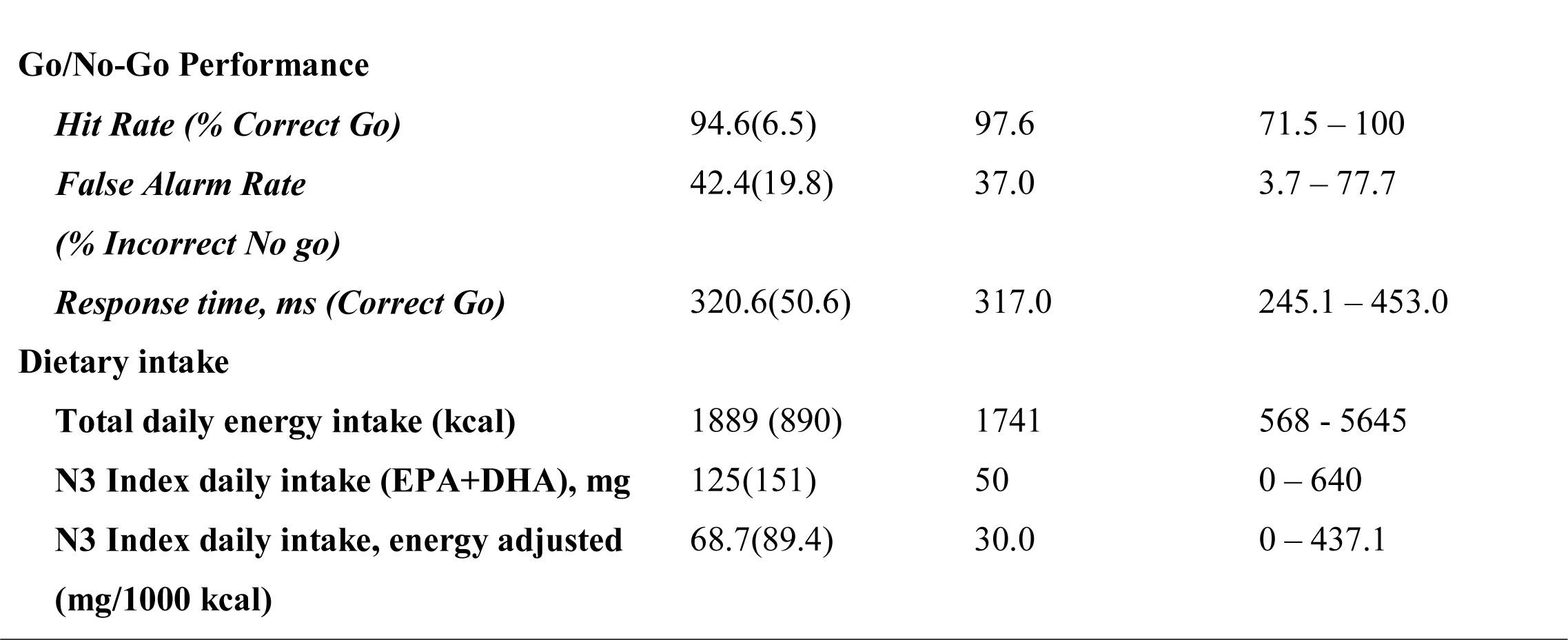
Participant characteristics

|  | Central Tendency |  |  |
| --- | --- | --- | --- |
|  | Mean(SD) | Median | Range |
| **N** | 87 |  |  |
| **Sex (% Females)** | 45 females<br>(51.7%) |  |  |
| **Age** | 13.3(1.1) | 13.3 | 11.1 – 16.1 |
| **Intelligence [n=84]** | 110.9(14.7) | 111 | 71 – 138 |
| **Race and Ethnicity** |  |  |  |
| *Caucasian* | 52.9% |  |  |
| *African American* | 28.7% |  |  |
| *Hispanic or Latino* | 9.2% |  |  |
| *Other* | 9.2% |  |  |
| **Socioeconomic Status index (z-score)**<br>**[n=84]** | 0.115(0.967) | 0.371 | -2.589 – 1.511 |
| *Parental education (years, mean)* | 16.5(2.7) | 17 | 7 – 22 |
| *Household income (median)* | \$50,000 - \$74,999 | \$100,00-\$149,999 | <\$5,000 - >\$200,000 |
| **Pubertal development** | 2.4(0.7) | 2.4 | 1 – 3.8 |
| **BMI z-score** | 0.34(0.93) | 0.40 | -2.2 – 2.2 |
| *Percentile* | 59.8(28.2) | 66.6 | 1.5 – 98.7 |
| **BRIEF Inhibit subscale, Percentile**<br>**(t-scores analyzed)** | 59.0(23.7) | 58 | 23 – 99 |
| **Go/No-Go Performance** |  |  |  |
| *Hit Rate (% Correct Go)* | 94.6(6.5) | 97.6 | 71.5 – 100 |
| *False Alarm Rate* | 42.4(19.8) | 37.0 | 3.7 – 77.7 |
| *(% Incorrect No go)* |  |  |  |
| *Response time, ms (Correct Go)* | 320.6(50.6) | 317.0 | 245.1 – 453.0 |
| **Dietary intake** |  |  |  |
| **Total daily energy intake (kcal)** | 1889 (890) | 1741 | 568 - 5645 |
| **N3 Index daily intake (EPA+DHA), mg** | 125(151) | 50 | 0 – 640 |
| **N3 Index daily intake, energy adjusted (mg/1000 kcal)** | 68.7(89.4) | 30.0 | 0 – 437.1 |

**Table 2.** MNI Coordinates of local maxima for activation during successful inhibitions (Correct No-Go>Incorrect No-Go) inversely associated with dietary N3 Index intake (cluster defining thresholdke=10, uncorrectedp=0.001, df 85). *Cluster survivingFWE correction atp<0.05 denoted.

| Region | *x* | *y* | *z* | Max *t* | Volume (mm3) |
| --- | --- | --- | --- | --- | --- |
| Inferior frontal gyrus, left | -42 | 2 | 24 | 3.47 | 12 |
| Insula, right | 46 | -2 | -2 | 4.05 | 25 |
| Insula, left | -40 | -2 | 6 | 3.87 | 30 |
|  | -35 | 0 | -2 | 3.57 |  |
| Anterior cingulate, left | -14 | 42 | -6 | 3.51 | 14 |
| Cingulate gyrus, right<br>[BA 32/24]* | 8 | 22 | 30 | 4.60 | 307* |
|  | 2 | 26 | 22 | 3.88 |  |

Age was unrelated to any behavioral variable (BRIEF Inhibit *t*-score [*r_s_*<-0.001, p=0.998, n=86], false alarms [*r_s_*=−0.142, p=0.190, n=87], and processing speed [*r_s_*=−0.130, p=0.231, n=87]). Though SES was unrelated to task performance (false alarms [*r_s_*=−0.136, p=0.218, n=84] and processing speed [*r_s_*=0.109, p=0.323, n=84]), greater family SES was associated with better caregiver-rated inhibitory control (BRIEF Inhibit *t*-score [*r_s_*=−0.311, p=0.004, n=83]).

### BRIEF inhibit subscale (parental report)

Mean and median percentile scores for this sample were 59% and 58%, respectively, indicating that the current sample is marginally more impulsive than peers of the same gender and age. Greater energy-adjusted intake of N3 Index was associated with better ability to inhibit behavior as rated by caregivers (BRIEF Inhibit subscale *t*-score) (*r_s_*=−0.257, p=0.017; n=86) (Figure 1).

**Figure 1.**
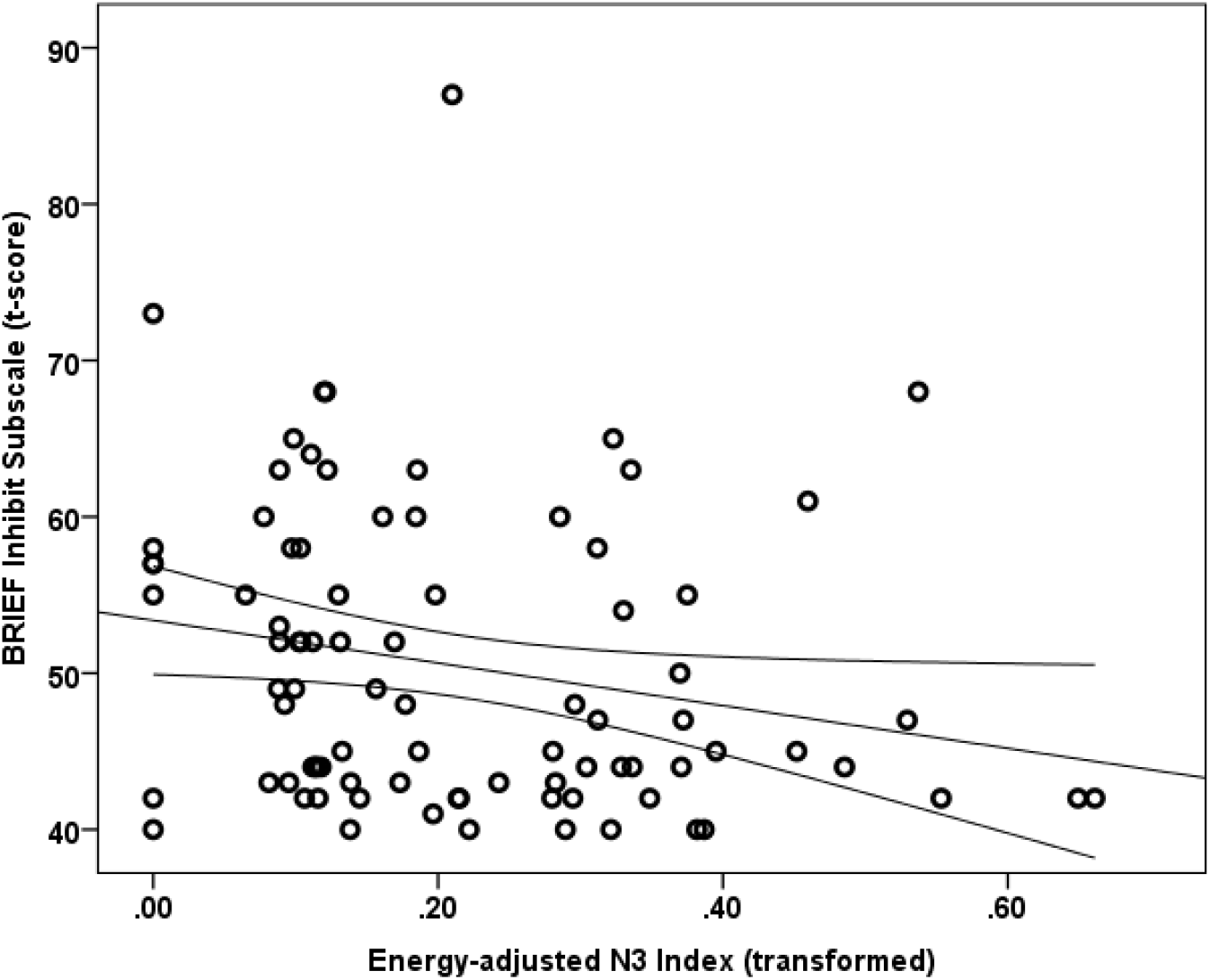
Energy adjusted N3 index intake is inversely related to response inhibition as rated by caregivers based on the BRIEF Inhibit subscale t-score (r_s_=−0.257, p=0.017).

A multiple linear regression was calculated using rank-transformed data to predict the Inhibit subscale ratings based on N3 Index and SES Index. Age was not a significant correlate of the Inhibit subscale score and thus was not included in the model. A significant regression equation was found (F(2, 80)=8.272, p=0.001), with an R^2^ of 0.171 (adjusted R Square 0.151). Standardized coefficients (β) were −0.274 (*t*=−2.678, p=0.009) for N3 Index and −0.281 (*t*=−2.743, p=0.008) for SES Index, indicating both SES index and energy adjusted N3 Index were significant predictors of BRIEF Inhibit subscale scores.

### Go/No-Go performance

Median false alarm rate (Incorrect No-Go) was 37%. The relationship between false alarm rate and energy-adjusted N3 Index did not reach statistical significance (*r_s_*=−0.194, p=0.072, n=87). Median reaction time (Correct Go’s) was 317 milliseconds. Reaction time was not significantly related to energy-adjusted N3 Index (*r_s_*= 0.157, p=0.145, n=87).

### PFC activity during successful response inhibition

Within the PFC, voxel-wise regression analysis revealed that activation during successful inhibitions (Correct No-Go>Correct Go) was inversely associated with energy-adjusted N3 Index in five clusters (Table 4). Figure 2 shows activation in the dorsal anterior cingulate cortex (dACC; 307 voxels; peak MNI 8, 22, 30; max *t* 4.60), which survived correction for multiple comparison (cluster level FWE, p=0.004; cluster level FDR p=0.014). There were no clusters where activation was positively correlated with N3 index. Beta weights were extracted from this cluster using MarsBar and plotted against energy-adjusted N3 Index for visualization purposes and to examine heterogeneity of activation (Figure 3).

**Figure 2.**
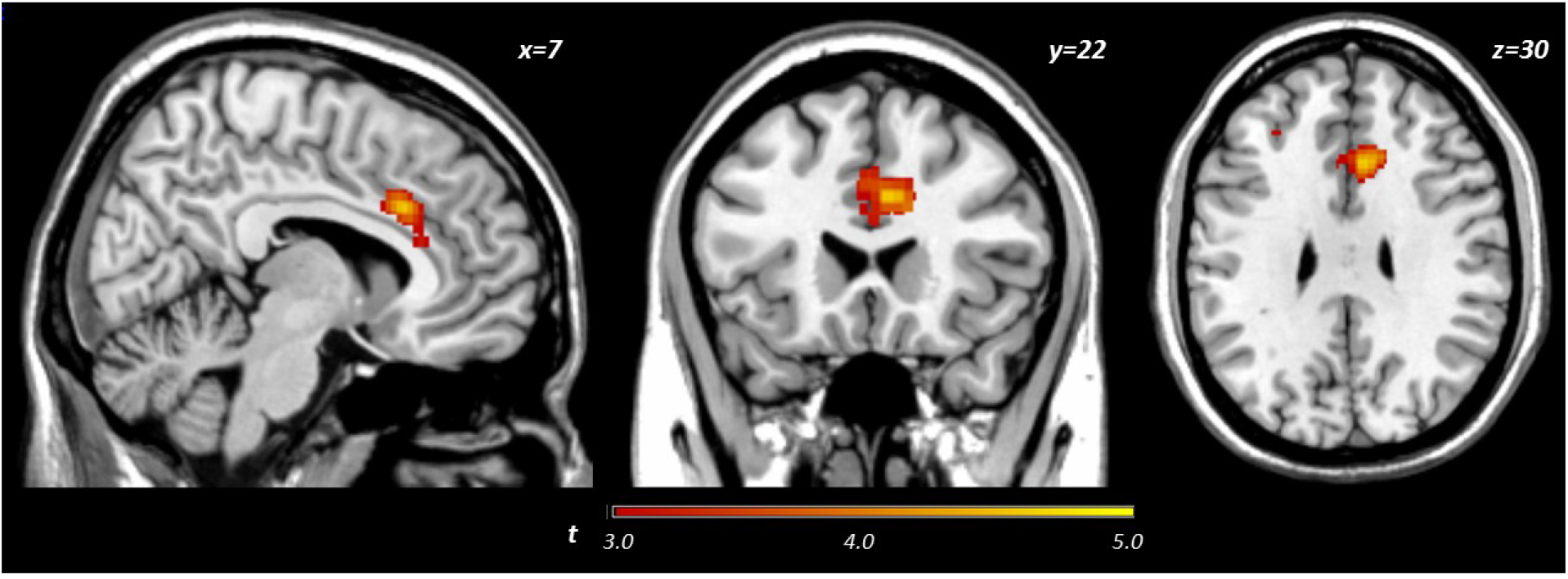
Activation during successful inhibitions inversely related to dietary N3 index.

**Figure 3.**
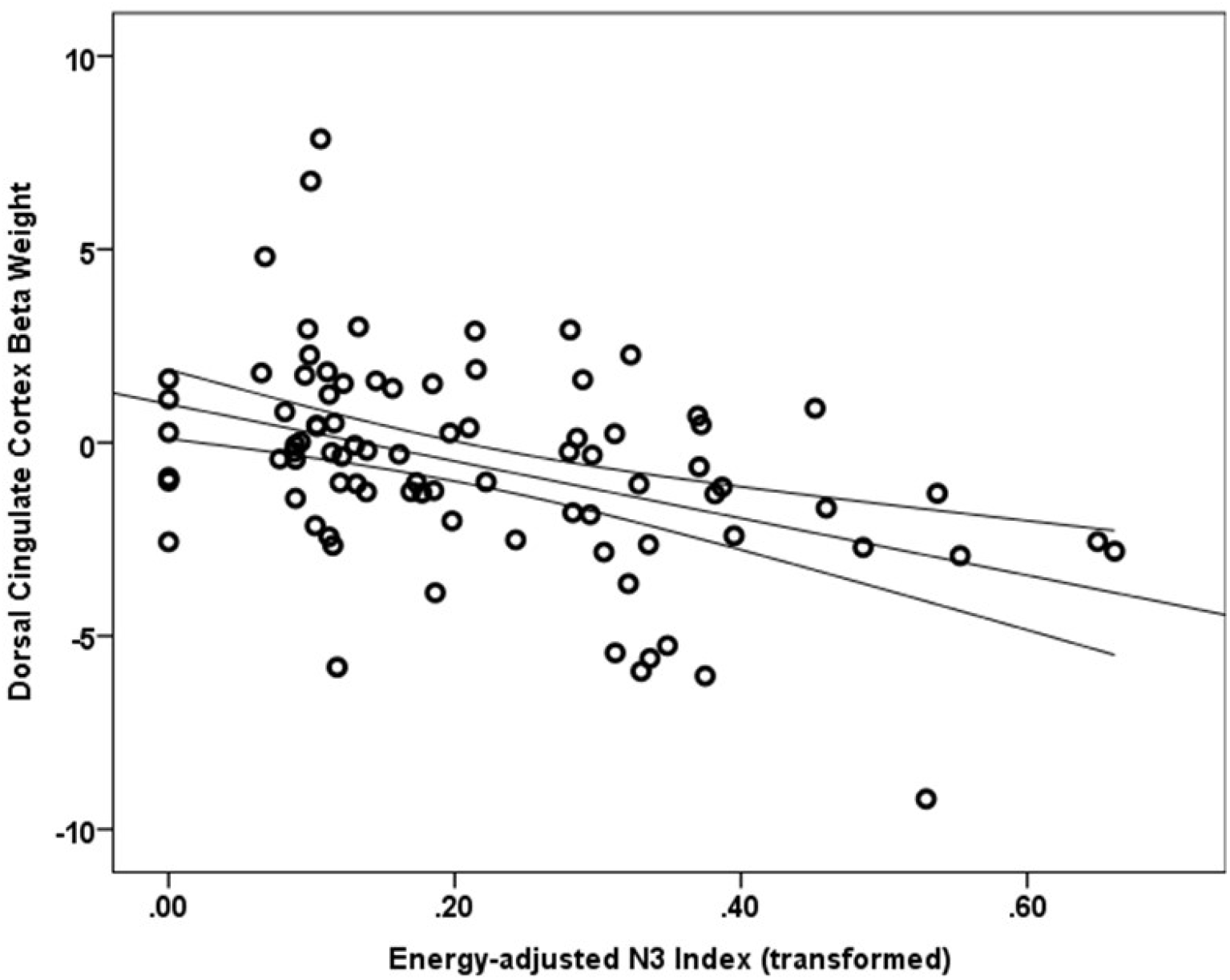
Relationship between dACC activation (regression coefficient beta weights) and energy-adjusted N3 Index intake.

## Discussion

The present study examined the extent to which dietary omega-3 fatty acids contribute to PFC function during successful impulse control in typically developing adolescents. We found that response inhibition during a simple Go/No-Go task was not significantly associated with reported energy-adjusted N3 Index. However, caregiver ratings of their child’s general ability to inhibit impulses were significantly inversely related to N3 intake; adolescents reporting lower N3 intake were rated as less able to control impulses in real-world situations. Furthermore, we found significant inverse relationship between energy-adjusted N3 Index and activity in the dACC during successful response inhibition, such that adolescents with lower omega-3 intake exhibit hyper-activation in the dACC in order to achieve similar behavioral performance as their peers reporting higher omega-3 intake. To our knowledge, this is the first study to correlate dietary long chain omega-3 fatty acids with impulse control and prefrontal function in adolescent boys and girls. Together with the extant comparable literature (reviewed below), the current findings suggest that long-chain omega-3 fatty acids may be particularly relevant to function of medial prefrontal cortex, particularly the dorsal region of the anterior cingulate cortex.

The PFC exerts top-down control over behavior and the anterior cingulate is putatively involved in, among many cognitive functions, performance monitoring during situations where the chance of error is high (Carter et al., 1998) or where there is need for heightened vigilance because conflicting responses are possible (Brown & Braver, 2005). Indeed, in adults, Go/No-Go task engagement recruits a network of regions including the dACC (Ogg et al., 2008). Among task events, successful inhibitions specifically recruit a network of regions including the rostral portion of superior medial frontal cortex (Simmonds et al., 2008). Though Simmonds et al. (2008) identified recruitment of a region slightly more dorsal than that reported in the present study (pre-supplementary motor area), the slight differences in activation coordinates may be partly due to inclusion of adults and contrasts examined (Correct No-Go > baseline versus Correct No-Go > Incorrect No-Go in the present study). Nevertheless, dACC function during error processing has been found to be critical to improvements in inhibitory control observed over development (Ordaz et al., 2013).

In the present study, consuming lower amounts of long chain omega-3 fatty acids was related to greater activity in the dACC during impulse control, but unrelated to task performance (rate of successful inhibitions). This result may suggest that for adolescents with lower omega-3 intake, greater dACC neural activity is required to accomplish the same level of inhibitory control as their high omega-3 intake peers. Omega-3 fatty acids are involved in functions at the cellular level that may represent a potential mechanism for the cortical inefficiency attributed to low N3 adolescents here. Interestingly, omega-3 deficiency has been shown to impair cortical glucose transport and utilization (Pifferi et al., 2005; Ximenes da Silva et al., 2002) and boys with lower omega-3 status exhibit indices of metabolic dysfunction in the ACC compared to their higher omega-3 peers (McNamara et al., 2013). Furthermore, omega-3 supplementation resulted in increased frontocortical efficiency in a rodent ADHD model (Liso Navarro et al., 2014). Relatedly, dACC activity attenuates over the course of engagement with the task (Tana, Montin, Cerutti, & Bianchi, 2010). Thus, it is possible that, in adolescents reporting a lower N3 Index, the cingulate experiences inefficient metabolism, or shows protracted habituation to the effort level required to perform the task, potentially signifying decreased efficiency. Additionally, given that DHA restriction leads to impaired pruning of superfluous axonal connections (de Velasco et al., 2012), it is possible that adolescents with lower N3 Index have greater activity in the ACC because this network has not yet undergone pruning of superfluous synapses, a concept proposed by Berl, Vaidya, & Gaillard (2006). Thus, inefficiency in metabolism and/or impaired cortical pruning may contribute to the greater ACC activity, which compensates for lower intake of long-chain omega-3 fatty acids.

The present results confirm previous reports of a relationship between omega-3 fatty acid status and the anterior cingulate (Almeida et al., 2017; Conklin, Gianaros, et al., 2007; McNamara et al., 2013). Further, since the current study includes both boys and girls, the present results extend the association to adolescent girls’ diet, dACC function and caregiver ratings of impulse control. It is notable that the current study also found a relationship between dietary N3 Index and cingulum activation without an *a priori* cingulate ROI. This confirms previous reports utilizing ROIs and suggests that midline structures may be sensitive to omega-3 levels (Almeida et al., 2017; Conklin, Gianaros, et al., 2007; McNamara et al., 2013). It should be noted that the region of interest for both McNamara et al. (2013) and Almeda et al. (2016) was more rostral and anterior to the cluster observed after a voxel-wise PFC analysis in the current study. Location of their ROI and other slight methodological differences may explain why we report a task-related difference while Almeida et al. (2016) did not observe a difference in ACC BOLD signal during task blocks requiring response vigilance/inhibition between boys with low and high blood omega-3 levels. Also, in contrast to Almeida et al. (2016) the current study distinguished between successful and unsuccessful trials in a hybrid design, rather than a block design, specifically to examine activation associated with successful response inhibition rather than attention *per se.* Additionally, the current study examined early adolescents (age 11-16 years) versus children (8-10 years), and included both males and females, where both developmental stage and sex have been demonstrated to have influence on developmental status/trajectory of BOLD signal (Ordaz et al., 2013) and neuroanatomical development (Giedd et al., 1999).

Consistent with our hypothesis, greater reported intake of omega-3 fatty acids was associated with better inhibitory control as rated by caregivers. Given that the BRIEF is not subject to adolescents’ self-report bias, our finding is notable in that it lends a degree of external validity to our results. Others have reported associations between omega-3 fatty acid status and selfreported impulsivity in adults (Conklin, Harris, et al., 2007) and in children (Gispert-llaurado et al., 2016). Together with previous studies, the present results support a role for omega-3 fatty acid status in generalized impulse regulation.

Contrary to our hypothesis, however, response inhibition as measured by ability to inhibit prepotent responses on the Go/No-Go task was unrelated to dietary N3 Index. Disparity between BRIEF subscales and presumably related task performance has been reported previously (McAuley, Chen, Goos, Schachar, & Crosbie, 2010). It is possible that assessing response inhibition using a task in a laboratory setting is a more circumscribed measure of inhibitory control, compared to caregiver report on general inhibitory control ability in real-world settings over the six months prior to the study visit. As acknowledged by Aron (2011), “the stopping of motor responses, no matter how sophisticated the model, will only be relevant for impulse control some of the time” (Aron, 2011). Consistent with our study, McNamara and colleagues also did not find differences between high and low DHA groups on false alarm rate using a similar task measuring response inhibition and sustained attention in boys (McNamara et al., 2010). Given that others have observed improved Go/No-Go performance (accuracy and response time) with supplementation in adults (Fontani et al., 2005), and have seen associations between (posterior) cingulate activation and performance during difficult but not easy task conditions (Boespflug, McNamara, Eliassen, Schidler, & Krikorian, 2016), it is possible that the intake levels reported in this study are too low and/or the current task is too easy (median hit rate 97.6%; median successful response inhibition rate 55.6%) to detect N3 associations with performance. Additionally, given that only a small, but significant, portion of the variance in attention (processing speed and omission errors) was attributable to Omega-3 Index in a large (n=266) cohort of adolescents (van der Wurff et al., 2016), it is possible that the current sample size (n=87) was underpowered to detect an association with task metrics of motoric response inhibition.

One of the main limitations of the current study is that the food frequency questionnaire is dependent on recall of usual diet. While other methods (24-hour recall, quantitative 7-food records) may increase accuracy, they are often difficult to implement with larger samples and require greater resources in terms of study team time and financial commitment. The food frequency questionnaire is an accepted method for measuring intake of nutrients with very high day-to-day variability, and its output represents the respondent’s chronic/habitual nutrient intake over specified periods of time (e.g., preceding 12 months) rather than a reliable calculation of absolute values (Subar et al., 2001). It is worth noting that reported intake has been previously shown to correlate well with serum biomarkers (Kuratko & Salem, 2009; Sun, Ma, Campos, Hankinson, & Hu, 2007; Vandevijvere et al., 2012), which was also demonstrated in a subset of participants in the current study providing both food frequency data as well as whole blood for essential fatty acid analysis. Furthermore, a review of reported global intake of dietary DHA found that 12-19 year-olds consume 30-50 mg/day in the U.S. (Flock, Harris, & Kris-Etherton, 2013), which is comparable to our sample (median reported daily intake DHA 30mg, data not shown). Thus, the present results demonstrate the relative reliability and feasibility of assessing diet in a moderately sized neuroimaging cohort. Some strengths of the present study include the inclusion of both male and female adolescents and a relatively restricted age range to minimize the variance of brain development due to chronological age.

### Summary and Implications

Dietary N3 Index intake was significantly inversely related to both activity in the dACC during successfully inhibited trials and to a general measure of impulsivity. Our results also reveal that energy-adjusted N3 Index accounts for a similar amount of unique variance in caregivers’ ratings of adolescent inhibitory control as accounted for by socioeconomic status, an established environmental factor in brain structure and function (Johnson, Riis, & Noble, 2016).

Dietary intake of effective omega-3 fatty acids, compared to omega-6 fatty acids, which are a competitive substrate for metabolism, has been on the decline in the U.S. over the past century (Blasbalg et al., 2011). While there are no dietary reference intakes for long chain omega-3 fatty acids (EPA and DHA), the consensus among experts is that the current level of intake is far below that desired for optimal health (Flock et al., 2013). A general theme in development is that the central nervous system’s DHA requirements are heightened during rapid growth of specific tissues. For example, early in postnatal development the retina has increased requirements for DHA and deficiency during this period reliably results in poor visual acuity (Agostoni, 2008). It is conceivable that the PFC may be comparably sensitive to insufficiency of DHA though over the longer developmental period of adolescence. Post mortem examinations indicate that PFC DHA content increases until 18 years of age (Carver et al., 2001), suggesting protracted accumulation in cortex. Given that cortical DHA levels would be predicted to fall by 5% within a few months of an omega-3 deficient diet in adults (Umhau et al., 2008), the metabolic needs of the developing adolescent brain may render it more sensitive to DHA levels. Thus, reduced DHA intake during adolescence, may delay and/or limit proper development of the PFC during this critical period in development, potentially leading to negative long-term consequences related to executive function though this hypothesis remains to be formally tested. To our knowledge this is the first larger-scale neuroimaging study in a sample of typically developing male and female adolescents to report a relationship between N3 Index and the ability to inhibit responses, a key executive function, as well as associated neural activity. While not evidence of a causal relationship, taken together these results suggest that intake of long chain omega-3 fatty acids is related to caregiver perceptions of their adolescent’s ability to control impulses and function of the dACC, a prefrontal region implicated in a number of executive functions including monitoring errors and performance. Unlike other comparable contributors like socioeconomic status *per se,* diet is a factor that may be more easily modified in the service of catalyzing morphological and functional neurodevelopment, specifically to increase behavioral self-control, which ultimately may have an impact on preventing maladaptive outcomes.

